# Relationship between protein thermodynamic constraints and variation of evolutionary rates among sites

**DOI:** 10.1101/009423

**Authors:** Julian Echave, Eleisha L. Jackson, Claus O. Wilke

**Affiliations:** Escuela de Ciencia y Tecnología, Universidad Nacional de San Martín, Martín de Irigoyen 3100, 1650 San Martín, Buenos Aires, Argentina; Department of Integrative Biology, Center for Computational Biology and Bioinformatics, and Institute for Cellular and Molecular Biology, The University of Texas at Austin, Austin, TX, USA

**Keywords:** protein evolution, rate variation among sites, biophysical model, thermodynamics, stability, stress

## Abstract

Evolutionary-rate variation among sites within proteins depends on functional and biophysical properties that constrain protein evolution. It is generally accepted that proteins must be able to fold stably in order to function. However, the relationship between stability constraints and among-sites rate variation is not well understood. Here, we present a biophysical model that links the thermodynamic stability changes due to mutations at sites in proteins (ΔΔ*G*) to the rate at which mutations accumulate at those sites over evolutionary time. We find that such a “stability model” generally performs well, displaying correlations between predicted and empirically observed rates of up to 0.75 for some proteins. We further find that our model has comparable predictive power as does an alternative, recently proposed “stress model” that explains evolutionary-rate variation among sites in terms of the excess energy needed for mutants to adopt the correct active structure (ΔΔ*G*\*). The two models make distinct predictions, though, and for some proteins the stability model outperforms the stress model and vice versa. We conclude that both stability and stress constrain site-specific sequence evolution in proteins.

## 1. Introduction

The evolution of protein-coding genes is shaped by functional and biophysical constraints on the expressed proteins (Pal et al. 2006, Thorne 2007, Worth et al. 2009, Wilke & Drummond 2010, Grahnen et al. 2011, Liberles et al. 2012). These constraints create patterns of rate variation *among* and *within* proteins. Among proteins, the primary determinant of rate variation is gene expression level (Drummond & Wilke 2008), though many other factors have been identified that also contribute to rate variation (Lemos et al. 2005, Xia et al. 2009, Liao et al. 2010, Pang et al. 2010). Within proteins, the primary determinants of rate variation seem to be linked to geometrical properties of the folded protein, in particular the Relative Solvent Accessibility (RSA) (Bustamante et al. 2000, Dean et al. 2002, Franzosa & Xia 2009, Ramsey et al. 2011, Shahmoradi et al. 2014) and the Local Packing Density (LPD) (Liao et al. 2005, Franzosa & Xia 2009, Yeh et al. 2014a, Yeh et al. 2014b) of sites in the three-dimensional protein structure.

To develop a mechanistic understanding of the causes that link geometrical properties, such as RSA and LPD, with site-specific rates of evolution, we need to develop explicit models of protein evolution. For example, recently a mechanistic “stress model” was proposed to explain the LPD-rate relationship (Huang et al. 2014). According to this stress model, LPD is a proxy of the stress energy ΔΔ*G*\*, a thermodynamic quantity that is a measure of the excess free energy needed for a folded mutant protein to adopt the correct active conformation. The stress model considers the effect of the stress free energy difference ΔΔ*G*\* but not that of possible mutational changes on global stability ΔΔ*G*. However, most proteins will function properly if they have folded *stably* into the correct conformation. To what extent stability constraints shape site-specific sequence evolution is not known.

Recent work has shown that describing protein evolution from the perspective of thermodynamic stability provides a wealth of insight into important aspects of protein evolution, such as the evolution of mutational robustness (Bloom et al. 2007), the origin of epistatic interactions (Bershtein et al. 2006, Gong et al. 2013), lethal mutagenesis (Chen & Shakhnovich 2009), determinants of evolutionary rate at protein level (Drummond & Wilke 2008, Serohijos et al. 2012), the evolution of novel function (Bloom et al. 2006, Tokuriki et al. 2008), and the expected equilibrium distributions of stability and the explanation of marginal stability (Taverna & Goldstein 2002, Goldstein 2011, Wylie & Shakhnovich 2011). Moreover, some studies suggest that ΔΔ*G*-based models are useful to study site-specific constraints. For example, Bloom & Glassman (2009) have shown that changes in stability upon mutation (ΔΔ*G* values) are intimately linked to the patterns of amino-acid substitutions observed over evolutionary divergence, to the extent that ΔΔ*G* values can actually be inferred with accuracy comparable to state-of-the art structure-based methods solely from an alignment of diverged protein sequences. More recently, Arenas et al. (2013) have used stability-based models to predict site-specific amino acid distributions. Despite the recognized importance of folding stability, stability-based models have not been used to predict the variation of evolutionary rates among sites.

Here, we investigate the relationship between mutational changes of stability and the site-dependency of rates of substitution. Following Bloom & Glassman (2009), we derived a neutral “stability model” of evolution which relates the ΔΔ*G*s due to mutations at a site with the site’s rate of substitution. For a diverse set of more than 200 enzymes, we compare the predicted rates with empirical rates (inferred from multiple sequence alignments) and with predictions of the stress model. The ΔΔ*G*-based and ΔΔ*G*\*-based predictions have on average similar correlations with empirical rates. However, the two models make significant independent contributions, which suggests that both stability and stress mould sequence divergence.

## 2. Stability Model: ΔΔ*G*-based rates

Our stability model is based on earlier work by Bloom and coworkers (Bloom & Glassman 2009, Bloom et al. 2005). The core idea of Bloom’s model is that there is a stability threshold Δ*G*^threshold^ such that all proteins more stable than the threshold are neutral (i.e. have all the same fitness) whereas all proteins less stable than the threshold are inviable (have fitness = 0). Thus, if Δ*G* is the stability of a protein, then its fitness *f* (Δ*G*) is assumed to be:

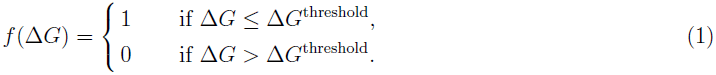

It is convenient to define

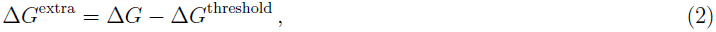

so that

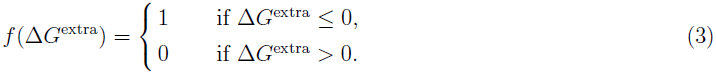

We further assume that the mutational effect on stability of a mutation *i → j* at site *k* is independent of the sequence background. We refer to this stability change as 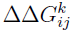. Because of the assumption of sequence independence, the stability difference between two sequences can be written as

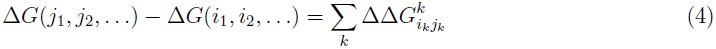

where *i*_1_ *i*_2,_… and *j*_1_, *j*_2_,… represent the amino acids of the two sequences, respectively. While this assumption cannot strictly be true, in practice it has worked well in several applications (e.g. Bloom et al. 2005, Bloom & Glassman 2009). The assumption is further supported by the observation that mutational effects on stability are frequently additive (Wells 1990, Serrano et al. 1993, Zhang et al. 1995) and tend to be conserved during evolution (Ashenberg et al. 2013).

Next we describe the evolutionary process. Throughout this work, we assume that the product of the protein-wide mutation rate *μ* and the effective population size *N*_*e*_ is small, *μN*_*e*_ ≪ 1. As a consequence, our populations are monomorphic, and we only have to track the evolution of a single representative sequence over time. We further assume that at most a single mutation arises at each time step.

The probability that a substitution *i* → *j* occurs at site *k* in a single time step, 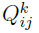 can be written as the product of the probability that the mutation *i* → *j* occurs, *M*_*ij*_, and the probability it goes to fixation

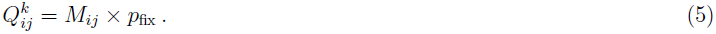

Here, we have assumed that all sites experience the same mutational process, so that *M*_*ij*_ does not depend on *k.* Note that *M*_*ij*_ scales with the effective population size *N*_*e*_, since all sequences in the population may mutate in one time step, and *p*_fix_ scales with 1/*N*_*e*_, because we are modeling the case of neutral evolution (Eq. 3). Thus *N*_*e*_ cancels, and we can set it equal to 1 without loss of generality.

Under the assumption of neutral evolution, the fixation probability is either one or zero, depending on whether the mutation keeps the extra stability in the negative or not. Because we have previously assumed that stability effects are independent of the sequence background (Eq. 4), they are fully specified by *i*, *j*, and *k.* (In other words, a mutation from *i* to *j* at site *k* always has the same stability effect 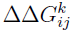.) However, the extra stability after the mutation, 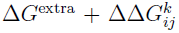, depends on the sequence background through the value of Δ*G*^extra^ before the mutation. From Eq. 3 we find the conditional fixation probability

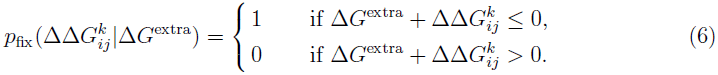

If 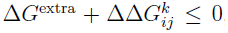 then the mutated protein is viable, and hence it fixes with probability 1. (Recall that we set *N*_*e*_ = 1.) By contrast, if 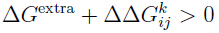, then the mutated protein is not viable and will not fix.

To proceed, we could write down a Markov process that keeps track of the extra stability at all time points (Bloom et al. 2007, Raval 2007). Instead, here we employ the “mean field” approximation of Bloom & Glassman (2009), in which we assume that Δ*G*^extra^ before mutation is drawn randomly from the steady-state distribution of Δ*G*^extra^ values, *p*_0_(Δ*G*^extra^), so that we can write the unconditional fixation probability as

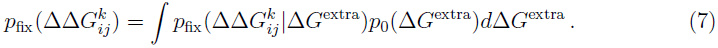

For *p*_0_(Δ*G*^extra^), Bloom & Glassman (2009) make the ansatz that it has an exponential probability-density function *p*_0_(Δ*G*^extra^):

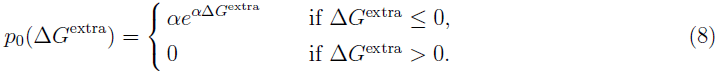

where *α >* 0 is a free parameter. This form cannot be derived from first principles, but it is justified by visual inspection of the probability density functions obtained under simulations (Bloom et al. 2007) (but see Wylie & Shakhnovich 2011).

Inserting Eq. 6 and Eq. 8 into Eq. 7, we obtain

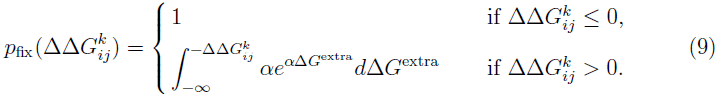

After taking the integral, we find

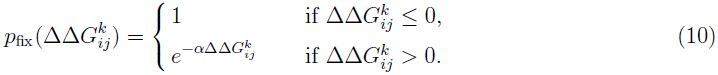

The stability model is completely specified by Eq. 5 and Eq. 10.

Next we consider the calculation of site-specific substitution rates. The substitution process at site *k* is described by a rate matrix **Q**^*k*^ with elements

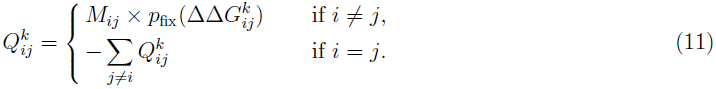

The stationary distribution 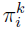 of the substitution process is given by the left null eigenvector of **Q**^*k*^, normalized such that 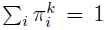. The rate of substitution at site *k*, 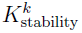, follows as

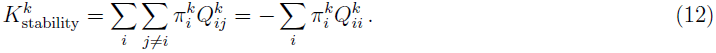

The subscript “stability” emphasizes that this rate estimate is calculated using the stability model.

In the case of symmetric mutations, *M*_*ji*_ = *M*_*ij*_, the equilibrium frequencies can be expressed as

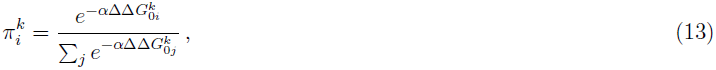

where 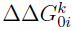 is the stability change relative to an arbitrarily chosen reference amino acid at site *k.* In the limit of unbiased mutations, *M*_*ij*_ = const for *i ≠ j*, the rate can be simplified to

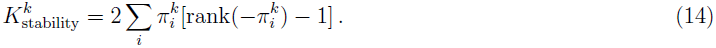

Here, rank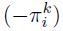 represents the rank order of 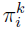, from largest to smallest. (The advantage of using Eq. 14 instead of Eq. 12 is that the latter contains a double-sum and hence is slower to evaluate.)

## 3. The Stress Model: ΔΔ*G*\*-based rates

The stability model is based on the assumption that fitness depends on whether the protein is stable enough to fold, so that the probability of fixation of a mutation will depend on the difference of folding free energy between the mutant and the wild-type, each in their respective equilibrium conformations. A different mechanistic model, the “stress model,” was recently derived based on the idea that, to be viable, a mutant must not only be stable, it must also be able to adopt a correct active conformation (Huang et al. 2014). Following this idea, the fixation probability of a mutant was modeled as the mutant’s probability of adopting the active conformation. According to this model, the rate of substitution for site *k* is

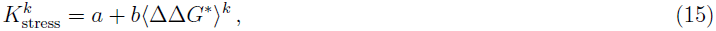

where ΔΔ*G*\* = Δ*G*_mutant_(**r**_active_) − Δ*G*_wt_(**r**_active_) is the free energy difference between mutant and wild-type when both adopt the active conformation and 〈ΔΔ*G*\*〉^*k*^ is its average over random mutations at site *k.* Since in general the active conformation will not necessarily be the relaxed equilibrium conformation, ΔΔ*G*\* represents the energy needed to stress the protein into adopting the right active conformation.

Further assuming that the active conformation is the wild type’s equilibrium conformation and approximating the free energy landscape using the parameter-free Anisotropic Network Model of Yang et al. (2009), it can be shown that 〈ΔΔ*G\**〉^*k*^ ∝ WCN^*k*^, where 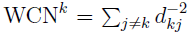 is the Weighted Contact Number introduced by Lin et al. (2008) and found to be among the best structural predictors of site-dependent evolutionary rates (Yeh et al. 2014a, Yeh et al. 2014b). Because of the proportionality between WCN^*k*^ and 〈ΔΔ*G*\*〉^*k*^, we can also write Eq. 15 as

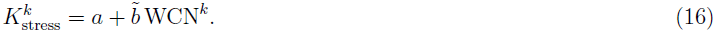

In practice, we obtain rates 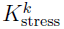 by calculating the WCN^*k*^ for each site *k* in a protein structure, fitting the linear expression 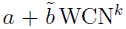 to a set of empirically estimated rates, and then using Eq. 16 to calculate a predicted rate at each site.

It is worthwhile to keep in mind that while the stability model takes into account whether the mutant is able to fold, the stress model takes into account the probability that the mutant adopts the right conformation. In principle both factors can affect fitness independently and therefore may both have an influence on substitution rates. If this is the case, both models are incomplete: the stability model does not consider the effect of possible conformational changes as long as the mutant is stable and the stress model takes stability for granted and considers only the destabilzation of the active structure.

## 4. Comparing the theoretical models with empirical data

### 4.1. Data set and calculation of empirical and predicted evolutionary rates

We tested our theory on the data set of Huang et al. (2014), which consists of 213 monomeric enzymes of known structure covering diverse structural and functional classes. Each structure is accompanied by up to 300 homologous sequences. In our analysis, we omitted four structures (1bbs, 1bs0, 1din, 1hpl) that had missing data at insertion sites. We aligned the homologous sequences for each structure with MAFFT (Multiple Alignment using Fast Fourier Transform) (Katoh et al. 2005, Katoh & Standley 2013). Using the resulting alignments as input, we inferred Maximum Likelihood phylogenetic trees with RAxML (Randomized Axelerated Maximum Likelihood), using the LG substitution matrix (named after Le and Gacuel) and the CAT model of rate heterogeneity (Stamatakis 2014).

For each structure, we then used the respective sequence alignment and phylogenetic tree to infer site-specific substitution rates with Rate4Site, using the empirical Bayesian method and the amino-acid Jukes-Cantor mutational model (aaJC) (Mayrose et al. 2004). The aaJC model poses equal probabilities for all amino-acid mutations, so that it is consistent with the theory presented in Section 2 and with the assumption of modeling amino-acid mutations as completely random perturbations made in the derivation of the stress model (Huang et al. 2014). Site-specific *relative* rates were obtained by dividing site-specific rates by their average over all sites of the protein, so that the mean relative rate of all sites in a protein was 1. In the following, we will refer to the rates inferred by Rate4Site as *empirical* rates, and will denote them by *K*_R4S_. We will refer to the rates calculated according to the stability model (*K*_stability_) or the stress model (*K*_stress_) as *predicted* rates. If necessary, we will distinguish between the predictions of the stability and stress model using the terms ΔΔ*G*-predicted rates and ΔΔ*G*\*-predicted rates, respectively.

We calculated ΔΔ*G* values with the program FoldX, following the default protocol (Guerois et al. 2002, Schymkowitz et al. 2005). Specifically, we first optimized the energy for each structure, using the RepairPDB method. We then calculated a 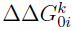 value for all possible 19 amino-acid substitutions at all sites in all proteins, using the PositionScan method, and considering the amino acid present in the PDB structure at each site as the reference amino acid at that site.

Rates predicted by the stability model were obtained using Eq. 13 and Eq. 14 either with *α* = 1 or with *α* chosen specifically for each protein. To determine the appropriate scale factor *α* for each protein, we maximized the correlation coefficient between the predicted site-specific rates as given by Eqs. 13 and 14 and the empirical site-specific rates as calculated by Rate4Site. To calculate the rates predicted by the stress model, we performed a linear fit between the site-dependent *K*_R4S_ and WCN for each protein, and then used Eq. 16 to calculate *K*_stress_ at each site.

All statistical analysis was carried out with R (R Core Team 2014). To fit the stability model to the data, we used the built-in function `optimize()` with default parameter settings. To fit the stress model to the data, we used the built-in function `lm()`. Correlation coefficients between predicted and empirical rates were calculated using `cor()` and partial correlations were obtained using the function `pcor.test()` of package `ppcor`.

All data and analysis scripts necessary to reproduce this work are available at: https://github.com/wilkelab/therm_constraints_rate_variation/.

### 4.2. Relationship between empirical and predicted evolutionary rates

We found that rates predicted by the stability model correlate significantly with the empirical rates. Correlation coefficients ranged between 0.25 and 0.75, with a median of 0.57 (Figure 1A). Scale values *α* fell between 0.52 and 2.63, with a median of 1.19. We further found that correlation coefficients and scale values were not correlated (*r* = 0.05, *P* = 0.47). To determine to what extent optimizing *α* for each protein affected the resulting correlation coefficients, we also calculated correlation coefficients with *α* = 1 for all proteins. We found that adjusting *α* made only a small difference, resulting on average in an increase in correlation coefficient of 0.007 (Figure 1A).

**Figure 1.**
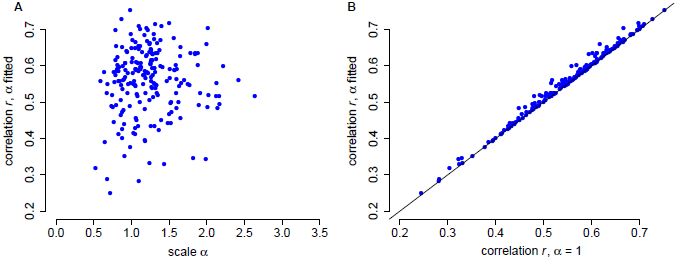
Correlations between rates predicted from ΔΔ*G* and rates inferred by Rate4Site. (A) Correlation coefficients vs. the fitted, protein-specific scales *α.* Each dot represents data for one protein. There is no relationship between the correlation coefficients and *α* (*r* = 0.10, *P* = 0.16). (B) Fitted *α* values provide only a small benefit over *α* = 1. Fitting *α* to each protein increases correlation coefficients, on average, by 0.007 (paired *t*-test, mean difference 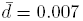, df = 208, *P <* 10^−10^).

We next investigated the functional relationship between empirical rates and rates predicted by the stability model. We pooled the data from all sites in all 209 proteins and calculated the joint distribution of the two rates. We also grouped sites into 20 bins of similar number of points using quantile breaks along the predicted rates axis. Figure 2 shows the joint distribution as well as the mean empirical rates and the 25% and 75% quantiles for each bin. The mean empirical rates fall nearly on top of the *x* = *y* line (which represents a perfect fit), with only a small amount of curvature around the mean predicted rate. The correlation between average empirical and predicted rates is *r* = 0.995, consistent with a very good linear fit. Despite the good fit of avarage rates, there is significant variation around *x* = *y,* as can be seen from the dispersion of the joint distribution around the *x* = *y* line and the error bars in Figure 2. The overall square correlation between ΔΔ*G*-predicted rates and empirical rates is *r*^2^ = 0.31, so that 69% of the variance of empirical rates is not explained by the stability model.

**Figure 2.**
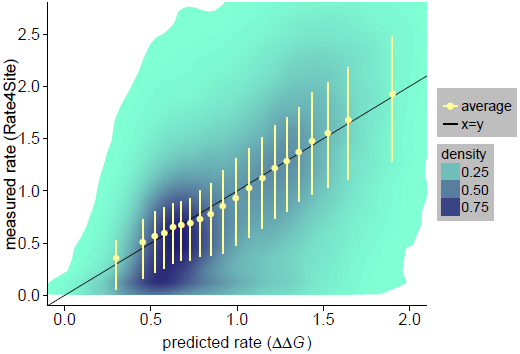
The relationship between rates predicted from ΔΔ*G* and rates inferred by Rate4Site is nearly linear. The joint distribution of empirical vs. predicted rates is shown using shaded areas. All sites were grouped into 20 bins of approximately equal number of sites using quantile breaks on the predicted rate axis. Yellow dots are the mean rates obtained by averaging over sites within a bin. Yellow error bars correspond to the 25% and 75% quantiles for each bin. Average empirical rates (yellow circles) are very close to the *x* = *y* line that corresponds to a perfect empirical-predicted fit (the correlation coeffient between mean empirical and predicted rates is 0.995). However, there is substantial variation around the mean trend, as can be seen from shaded areas and yellow error bars (correlation between non-averaged empirical and predicted rates is 0.558).

Next, we compared the predictions of the stability model with those from the stress model (Huang et al. 2014), which describes site-specific evolutionary rates in terms of the increased stress that results in the protein’s active conformation due to mutation (ΔΔ*G*\*). In a protein-by-protein comparison, the stability model is somewhat better (dots above the *x* = *y* line in Figure 3) for 127 of the 209 proteins, a proportion significantly larger than 50% (binomial test: 61%, *P* = 0.002). When considering all sites together, the two models perform comparably. The correlations between empirical and predicted rates for all sites are 0.56 with ΔΔ*G*-based predictions and 0.55 with ΔΔ*G*\*-based predictions. However, even though the two models perform comparably on average, there is substantial variation around the mean trend (Figure 3). For some proteins, the ΔΔ*G* model clearly outperforms the stress model and vice versa. Also, considering all sites, the partial correlations between empirical rates and predicted rates for one model controling for the predictions of the other are 0.33 and 0.31 for the ΔΔ*G* model and the stress model, respectively. These values are large and highly significant (*P* ≪ 10^−3^), showing that the predictions of the two models are quite independent and may be accounting for different constraints.

**Figure 3.**
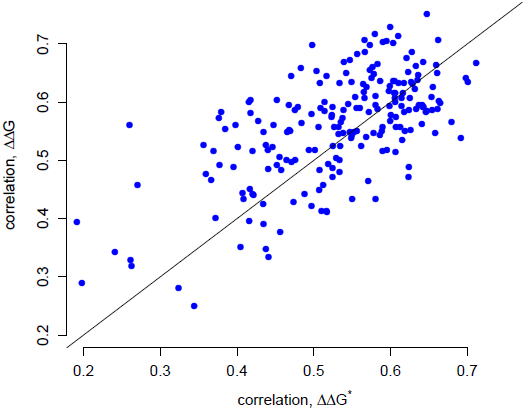
Correlations between rates inferred by Rate4Site and rates predicted by either the stress ΔΔ*G*\*-based model (shown along the *x* axis) or the stability ΔΔ*G-*based model (shown along the *y* axis). The correlation coefficients from the two models are significantly correlated (*r* = 0.64, *P <* 10^−10^). Correlations have similar magnitudes, with the ΔΔ*G*-based model giving slightly better results on average (paired *t*-test, mean difference 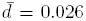, df = 208, *P <* 0.001). For 127 of the 209 proteins the stability model gives better correlations while for 82 of the 207 proteins the stress model gives better results.

The relative independence of stress and stability as determinants of site-specific evolutionary rates suggests that considering both factors should improve predictions. To verify this hypothesis, we fit empirical rates to a linear combination of rates predicted from ΔΔ*G*\* and ΔΔ*G*. Considering all sites of all proteins, the two-variable model results in a square correlation *R*^2^ = 0.38, approximately a 23% improvement over *R*^2^ = 0.31 of the stability model and *R*^2^ = 0.30 of the stress model. Both predictors in the two-variable model are highly significant (*P* < 10^−15^). These results further support the idea that stability and stress provide significant independent constraints to evolutionary divergence at site level.

All ΔΔ*G*-based predictions presented above used ΔΔ*G* values calculated by FoldX. It is possible that a different ΔΔ*G* predictor would yield substantially different results. In particular, even though FoldX is a state-of-the-art ΔΔ*G* predictor its predictions explain only 25% of the variance in measured ΔΔ*G* values (Potapov et al. 2009, Thiltgen & Goldstein 2012), indicating a substantial need for improved ΔΔ*G* prediction methods with higher accuracy. Therefore, we also asked to what extent our results depended on the method by which we calculated ΔΔ*G* values. We calculated a second set of ΔΔ*G* values, using the ddg_monomer application in Rosetta (Kellogg et al. 2011). Because this application runs approximately 500 times slower than FoldX, we could not run it on all proteins in our data set. Instead, we arbitrarily selected five proteins (PDB IDs 1bp2, 1lba, 1ljl, 1pyl, and 2acy) as a test case. We found that FoldX performs similarly or better than ddg_monomer (Figure 4). Thus, in our application here, we could not identify any major differences between predictions obtained from FoldX and those obtained from Rosetta ddg_monomer.

**Figure 4.**
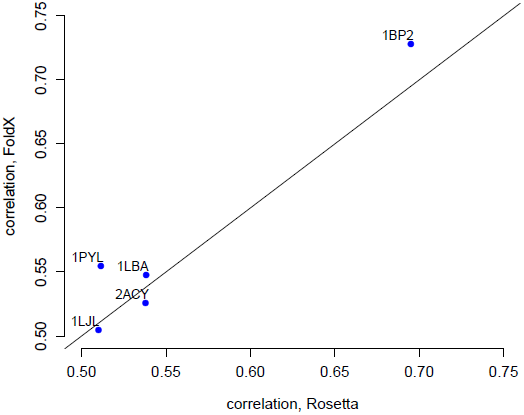
Rates predicted using ΔΔ*G* values obtained from FoldX perform as well as or better than the ones obtained from the ddg_monomer protocol in Rosetta. Shown are the correlation coefficients of measured rates with rates predicted using the stability model with FoldX ΔΔ*G* values (y axis) vs. Rosetta ΔΔ*G* values (x axis) for five proteins.

### 5. Conclusion

We have developed a biophysical model linking stability changes ΔΔ*G* due to mutations at individual sites in proteins to site-specific evolutionary rates. This stability model predicts site-specific rates in very good agreement with empirical rates. Indeed, the overall correlation between empirical rates and ΔΔ*G*-based predictions is similar to the correlation with the best structural determinant, the packing density measure WCN, which, according to a recent mechanistic stress model, is a measure of the local stress introduced by mutations into the active protein structure ΔΔ*G*\* (Yeh et al. 2014b, Huang et al. 2014). However, despite the similar performance, large partial correlations show that the two factors ΔΔ*G* and ΔΔ*G*\* result in largely independent predictions. Moreover, there are proteins for which the stability model performs significantly better than the stress model, while for other proteins the reverse is true. Consistently, a two-variable model that combines stability and stress significantly improves predictions. Therefore, both the overall stability ΔΔ*G* and the stress ΔΔ*G*\* seem to capture distinct thermodynamic constraints on protein evolution.

The stability model presented here is a neutral model in which mutations are either neutral or completely deleterius according to whether the mutant’s stability is above a certain threshold (Taverna & Goldstein 2002, Bloom et al. 2005, Bloom & Glassman 2009). A presumably more sophisticated model is based on posing a continuous dependence between fitness and Δ*G* (Tokuriki & Tawfik 2009, Chen & Shakhnovich 2009, Goldstein 2011, Wylie & Shakhnovich 2011). However, even though the continuous fitness models appear to be more realistic than the neutral stability-threshold models, in a recent study Arenas et al. (2013) found that the neutral model leads to better predictions of site-specific amino-acid distributions. This finding provides additional support for our choice of using a neutral ΔΔ*G*-based model. In future work, it will be worthwhile to explore the site-dependency of substitution rates using continuous fitness-stability models.

## Acknowledgements

We would like to thank Stephanie Spielman for help with setting up the evolutionary-rate calculations. J.E. is a researcher of CONICET. E.L.J, is funded by an NSF Graduate Research Fellowship, grant number DGE-1110007. C.O.W. is supported by NIH grant R01 GM088344, DTRA grant HDTRA1-12-C-0007, ARO grant W911NF-12-1-0390, and the BEACON Center for the Study of Evolution in Action (NSF Cooperative Agreement DBI-0939454). The Texas Advanced Computing Center (TACC) provided high-performance computing resources.

